# NLRP3 controls ATM activation in response to DNA damage

**DOI:** 10.1101/2020.05.12.087015

**Authors:** Mélanie Bodnar-Wachtel, Anne-Laure Huber, Julie Gorry, Sabine Hacot, Laetitia Gerossier, Baptiste Guey, Nadège Goutagny, Birke Bartosch, Elise Ballot, François Ghiringhelli, Bénédicte F. Py, Yohann Coute, Annabelle Ballesta, Sylvie Lantuejoul, Janet Hall, Virginie Petrilli

## Abstract

The DNA damage response (DDR) is essential to preserve genomic integrity and acts as a barrier to cancer. The ATM pathway orchestrates the cellular response to DNA double strand breaks (DSBs), and its attenuation is frequent during tumorigenesis. Here, we show that NLRP3, a Pattern Recognition Receptor known for its role in the inflammasome complex formation, interacts with the ATM kinase to control the early phase of DDR, independently of its inflammasome activity. NLRP3 down-regulation in human bronchial epithelial cells impairs ATM pathway activation as shown by an altered ATM substrate phosphorylation profile, and due to impaired p53 activation, confers resistance to acute genomic stress. Moreover, we found that NLRP3 is down-regulated in Non-Small Cell Lung Cancer (NSCLC) tissues and NLRP3 expression is correlated with patient overall survival. NLRP3 re-expression in NSCLC cells restores appropriate ATM signaling. Our findings identify a non-immune function for NLRP3 in genome integrity surveillance and strengthen the concept of a functional link between innate immunity and DNA damage sensing pathways.

## INTRODUCTION

Maintenance of genome integrity is crucial for cell survival. Toxic DNA double strand breaks (DSBs) can arise from both exogenous sources, for instance exposure to ionizing radiation (IR), or endogenous sources such as DNA replication stress. If they remain unrepaired or are incorrectly repaired they represent a major risk factor for genome instability, a condition known to favor tumorigenesis. One of the key proteins that orchestrates the rapid cellular response to DSBs is the Ataxia-Telangiectasia Mutated (ATM) kinase. The early molecular mechanism(s) leading to ATM activation upon DSB formation remain elusive. In resting cells, ATM is present as an inactive dimer. Once recruited to DSBs via the action of the MRE11-RAD50-NBS1 (MRN) complex, ATM autophosphorylates, monomerizes and initiates a vast cascade of post-translational modifications that are essential for the DNA Damage Response (DDR) ^1^. Phosphorylation of the histone variant H2AX on Ser139 (γH2AX) is one of the earliest events in the DDR and is crucial for an efficient recruitment of DNA repair proteins to strand breaks to form Ionizing Radiation Induced Foci (IRIF) ^2–5^. The scaffold protein MDC1 binds to γH2AX and recruits more ATM to the DNA lesion thus amplifying and maintaining the DNA damage signal ^6,7^. ATM also phosphorylates many downstream effector proteins including KAP1, p53 and CHK2, which induce effector mechanisms such as the activation of cell cycle checkpoints, apoptosis or senescence ^8^. The ATM pathway is tightly regulated and any dysregulation in these protection mechanisms facilitates the progression of cells towards malignancy.

NLRP3 belongs to the Nod-Like Receptor (NLR) family, a family of cytosolic Pattern Recognition Receptors (PRRs) involved in innate immunity ^9^. Upon sensing of Pathogen-Associated Molecular Patterns (PAMPs) or Damage-Associated Molecular Patterns (DAMPs) such as nigericin or ATP, respectively, NLRP3 triggers the assembly of a multi-protein complex, the inflammasome, the function of which is to control caspase-1 activation ^10–12^. Activated caspase-1 induces the maturation of the pro-inflammatory cytokines IL-1β and IL-18, and eventually pyroptosis of the cell ^13,14^. The NLRP3 inflammasome is mostly expressed in macrophages and dendritic cells where its biological functions have been well characterized. Whether NLRP3 exerts functions unrelated to immunity remains unknown ^15,16^. A previous study reported that in myeloid cells the NLRP3 inflammasome activity relies on the presence of ATM, suggesting a functional link between these two pathways ^17^. Here, we investigated whether NLRP3 controls the ATM pathway. We discovered that NLRP3 binding to ATM is instrumental to early ATM activation and to trigger apoptosis in response to DNA DSBs, and we report that NLRP3 expression is down-regulated in NSCLC.

## Materials and Methods

### Cell culture

HBEC3-KT, HCC15, HCC366, HCC4017 and HCC461 were obtained from J. Minna, NCI-H1703 (H1703), NCI-H292 (H292), NCI-H520 (H520), NCI-H661 (H661), NCI-H358 (H358), NCI-H1792 (H1792), NCI-H441 (H441) and SK-MES-1 from ATCC, and A549 from the PHE culture collection. HBEC3-KT cells were cultured in Keratinocyte-Serum Free Medium (Invitrogen) supplemented with 12.5 mg of Bovine Pituitary Extract (Invitrogen) and 100 ng of epidermal growth factor (Invitrogen). H1792, A549, HCC15, HCC366, HCC4017, HCC461 and H441 were cultured in RPMI medium, supplemented with 10% Fetal Bovine Serum (FBS) and 1% penicillin/streptomycin, H1703, H292, H520, H358, H661 in RPMI, 10% FBS, 1 mM sodium pyruvate, 1 mM HEPES and 1% penicillin/streptomycin and SK-MES-1 in RPMI, 10% FBS, 1 mM sodium pyruvate, 1 mM non-essential amino acids and 1 % penicillin/streptomycin (Invitrogen). *NLRP3* mutated cell lines are H661, H358, HCC15, HCC366 and HCC4017. HeLa cells were cultured in DMEM 4.5 g/L of glucose, 10% FCS and 1% penicillin/streptomycin. Treated cells received Etoposide (TEVA santé), HU (Sigma) or γ-ray treatment 24 h post-transfection. For inflammasome activation, cells were primed overnight with 10 μg/mL poly(I:C) (Invivogen) and treated with nigericin (10 μM, 6 h), or ATP (5 mM, 30 min) (SIGMA). IL-1β ELISA was purchased from BD. Z-VAD-fmk and z-YVAD-fmk were from Enzo Life Science.

### Mice

The NLRP3 flox mice were generated by the “clinique de la souris” Strasbourg. The exon 4 was flanked by 2 Lox-P sites in C57BL6 background. NLRP3^flox/flox^ mice were bred with Rosa26-Cre-ERT2 mice. Bone marrow-derived macrophages were generated as previously described ^39^ from +/+; Cre-ERT2 and Flox/Flox; Cre-ERT2 adult mice. To inactivate *Nlrp3* (*Nlrp3*^ΔΔ^), 4OHT (SIGMA) was added at the concentration of 0.5 μM from day 2 to day 7 of differentiation in both genotypes for control. BMDM were treated with 100 μM Eto as indicated.

### Cell transfection

HBEC3-KT were seeded at 1.5×10^5^ cells per well in 6-well plates and were transfected with non-targeting siRNA (SMART pool Non-targeting siRNA, Dharmacon) or NLRP3 siRNA (SMARTpool, Dharmacon) using HiPerfect transfection reagent (Qiagen) or INTERFERin (Polyplus transfection) following manufacturer’s instructions. HeLa cells were transfected with Lipofetamin2000™ (Invitrogen). H292 cells were transfected using PEI method. Vectors used pCR3-Flag-NLRP3 (FL, SHORT, PYD, NACHT, LRR), pcDNA3.1-Flag-His-ATM (Addgene 31985), pcDNA3.1-mCherry-NLRP3. shRNA were from Genecopoeia, hygromycin selection, *NLRP3:* forward: 5’-TAATACGACTCACTATAGGG-3’ Reverse: 5’-CTGGAATAGCTCAGAGGC-3’; control Forward: 5’-TAATACGACTCACTATAGGG-3’ Reverse: 5’-CTGGAATAGCTCAGAGGC-3’.

### Irradiation

Cells were irradiated using a 6-MeV γ-ray clinical irradiator (SL 15 Phillips) at the Léon Bérard Cancer Centre (Lyon, France) with a dose rate of 6 Gy.min^−1^ to obtain the required dose.

### ROS measurement

For intracellular ROS staining, HBEC3-KT cells were incubated with 1 μM of 2’-7’-dichlorofluorescin diacetate (CM-H2DCFDA; Invitrogen) for 30 min at 37°C. For a positive control, cells were pretreated with 5 μM of ATM inhibitor (ATMi) (KU55933; Selleckchem) for 5 h prior to staining. Stained cells were collected and analyzed on a BD FACSCalibur, and data were analyzed using the FlowJo software.

### Mathematical modeling

ATM dynamics was modeled using the following ordinary differential equation:

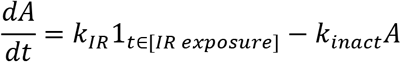

where A is the concentration of activated ATM molecules (expressed in number of foci/cell), *k_IR_* is the activation rate (in number of foci/cell. h^−1^), only present during irradiation, and *k_inact_* is the inactivation rate (in h^−1^). A was set at zero at t0. This model assumed that ATM molecules were in excess compared to the activated proportion.

The model is fitted to experimental data by estimating the optimal values for *k_IR_* and *k_inact_* for each condition-siCTL or siNLRP3-using a weighted least-square approach ^40^. For hypothesis A, i.e. NLRP3 enhances ATM activation, *k_IR_* is assumed to be different for both conditions whereas *k_inact_* is taken identical. For hypothesis B, i.e. NLRP3 inhibits ATM deactivation, *k_inact_* is assumed to vary between siCTL and siNLRP3 conditions and *k_IR_* is assumed to remain identical. All computations were done in Matlab (Mathworks, USA).

### Generation of NLRP3 stably expressing cells

The human *NLRP3* cDNA was inserted into the lentiviral vector pSLIKneo (addgene) containing a TET-ON promoter using the Gateway recombination system (Invitrogen). Sequences of the Gateway shuttle vectors are available upon request. Empty pSLIK vector (without the ccdB containing Gateway recombination cassette) was produced by partial digestion with Xba1 and Xho1 followed by religation. Viral particles were produced by transfecting HEK293T cells with the lentiviral vectors and a second generation packaging system. NCI-H292, NCI-H520 and A549 cells were either transduced with the empty pSLIK control vector or the NLRP3 containing vector. NLRP3 expression was induced by adding 0.5 μg/mL doxycycline (Takara Bio) to the cell culture medium.

### Western blotting

Cells were washed with PBS and detached by trypsinization. Cell pellets were lyzed in Laemmli buffer x2 (Tris HCl 0.5 M pH 6.8; 0.5 M DTT; 0.5% SDS) and protein concentrations were determined using the Bradford reagent (Biorad). Protein extracts were separated on SDS-PAGE (8 % or 15 % or 4-15% gradient (vol/vol)) gels. Gels were transferred onto nitrocellulose membranes (GE HealthCare and Biorad) for immunoblotting with the following antibodies: anti-NLRP3 (Cryo-2, 1:1000) and anti-caspase-1 (Bally-1, 1:1000) from Adipogen, anti-ASC (1:2000) from ENZO Life Science, anti-γH2AX (JBW301, 1:1000), anti-P-Ser15-p53 (1:1000) and anti-ATM Ser1981 (10H11.E12, 1:200) were from Millipore. Anti-P-KAP1Ser824 (1:1000), anti-KAP1 (1:1000) and anti-Nek7 (A302-684A) from Bethyl Laboratories, anti-p53 (clone DO7 1:2000) and anti-NOXA (114C307, 1:1000) from Santa Cruz, anti-ATM (#ab32420, 1/1000) from Abcam, anti-Flag (F7425 1/5000) from Sigma, anti-XPO2 (GTX102005 1/1000) and anti-IPO5 (GTX114515 1/1000) from Genetex and anti-actin (C4, 1:100,000) from MP Biomedical.

The Fiji and ImageLab programs were used for densitometric analysis of immunoblots, and the quantified results were normalized as indicated in the figure legends.

### Cell fractionation

HBEC3-KT were fractionated by adapting the method described by Hacot et al. ^41^. The MgCl_2_ concentration used for the hypotonic buffer was 0.75 mM. Equal amounts of proteins were run by SDS-PAGE.

### Immunofluorescence

Cells were plated onto sterile glass coverslips and fixed with PBS-PFA 4% (wt/vol) for 15 min at room temperature (RT) and washed twice in PBS. Cells were permeabilized with lysis buffer (sucrose 300 mM, MgCl_2_ 3 mM, Tris pH 7.0 20 mM, NaCl 50 mM, Triton X-100 0.5%) for 3-5 min at RT under slow agitation. The following antibodies were diluted in PBS-BSA 4% and applied to the coverslips for 40 min at 37°C: anti-γH2AX (JBW301, 1:800), P-ATM Ser1981 (10H11.E12, 1:200), and 53BP1 (1:500) from Millipore, and anti-MDC1 (1:200) from AbCam. For NLRP3 labeling, cells were fixed with PBS–PFA 4% (wt/vol) for 15 min at RT, washed twice in PBS and permeabilized with 1% triton X100. Anti-flag was diluted in saturation buffer (PBS, 1% BSA; NaCl 0.02%; Tween 20 0.5%; SVF 3%) and incubated on cells for 1 h. Cells were then incubated with Alexa-Fluor 488-conjugated anti-mouse or Alexa-Fluor 555-conjugated anti-rabbit (1:800; Life Technologies) for 20 min at 37°C and in Hoechst (500 ng/mL in PBS) for 10 min at RT. Fluorescence microscopy pictures were taken using a Nikon Eclipse Ni-E microscope, and confocal Zeiss LSM 780. The Fiji program was used to analyze fluorescence intensity.

### Live imaging

mCherry-NLRP3 transfected H292 were imaged using a confocal spinning disk inverted microscope (Leica, Yokogawa). Vital Hoechst was used at 0.5 μg/mL. The Fiji program was used to analyze fluorescence intensity.

### Co-immunoprecipitation

HeLa cells were transfected using Lipofectamin2000™ (Invitrogen) according to manufacturer’s protocol, and lyzed in the following buffer: Tris HCl 100 mM pH 8.0, 10 mM MgCl_2_, 90 mM NaCl, 0.1% Triton X-100, Complete® tablet (Roche). Immunoprecipitation was performed using M2-agarose beads (A2220 Sigma) overnight at 4°C.

### Proximity Ligation Assay

HBECT3-KT and HeLa cells were seeded onto glass coverslips and processed as described by the manufacturer’s protocol (Duolink® PLA Technology, Sigma). Antibodies used NLRP3 1/500 (ABF23, Millipore), ATM 1/500 (2C1, Abcam). Quantification was carried out using the macro published by Poulard et al ^42^.

### IL-1β Luminex Assay

IL-1β levels in cell supernatants were analyzed using the Magnetic Luminex Screening Assay according to the manufacturer’s protocol (R&DSystems). The samples were read using the Bioplex-200 from BioRad.

### Quantitative reverse transcription PCR

Cells were washed and detached by trypsinization. RNA was extracted using NucleoSpin^®^ RNA kit (Macherey Nagel). Five hundred nanograms to one microgram of RNA were reverse transcribed using SuperScript II reverse transcriptase and oligo(dT) primers (Life technologies) and RNAsin (Promega). cDNAs were quantified by real-time PCR using a SYBR^®^ Green PCR Master Mix (Applied Biosystems) on a ABI Prism^®^ 7000 (Applied Biosystems) or CFX Connect Real-Time system (BioRad). Sequences of the primers NLRP3 Forward 5’-GAAGAAAGATTACCGTAAGAAGTACAGAAA; Reverse 5’-CGTTTGTTGAGGCTCACACTCT; ESD 5’-TTAGATGGACAGTTAC TCCCTGATAA; Reverse 5’-GGTTGCAATGAAGTAGTAGCTATGAT; HPRT1 Forward 5’-CATTATGCTGAGGATTTGGAAAGG; Reverse 5’-TGTAGCCCTCTGTGTGCTCAAG; CBP Forward 5’-CGGCTGTTTAACTTCGCTTC; Reverse 5’-CACACGCCAAGAAACAGTGA. NOXA Forward 5’-GGAGATGCCTGGGAAGAAG; Reverse 5’-CCTGAGTTGAGTAGCACACTCG; PUMA Forward 5’-CCTGGAGGGTCCTGTACAATCTCAT; Reverse 5’-GTATGCTACATGGTGCAGAGAAAG; ACTIN Forward 5’-AGCACTGTGTTGGCGTACAG; Reverse 5’-TCCCTGGAGAAGAGCTACGA.

NLRP3 mRNA amounts in different NSCLC cell lines were normalized against *ESD* (Fig. 1B) or in human samples on *CPB* and *HPRT1* (Fig. 1D) mRNA levels or in HBEC-KT on *HPRT1* (Fig. 5C).

**Figure 1.**
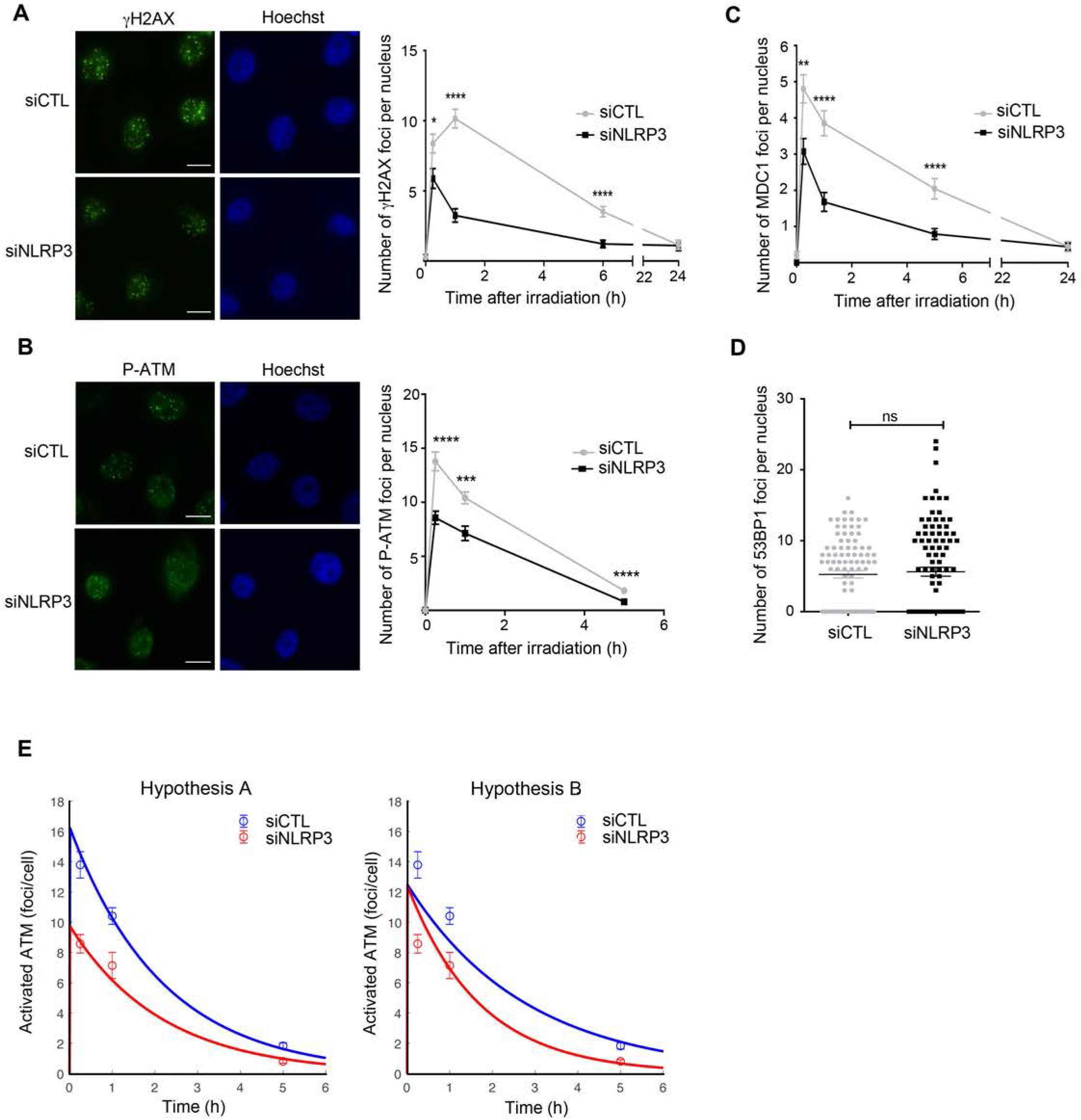
NLRP3 down-regulation impairs ATM-dependent IRIF formation and DNA damage signaling in response to DNA double strand breaks. (**A** to **D**) HBEC3-KT transfected with control (siCTL) or NLRP3 siRNA (siNLRP3) were treated with IR (2 Gy). The number of nuclear γH2AX (A), MDC1 (**B**), P-ATM (**C**), and 53BP1 (**D**) IRIF were determined by immunofluorescence at indicated time points. IF of the 1 h time point is shown (x60). Hoechst (blue) was used to stain nuclei. Representative of four (**A**, **C**) and two independent experiments (**B**, **D**) (64≤ n ≥148) Scale bars 10 μm. (**E**) Mathematical modeling of ATM and NLRP3 interactions. Two hypotheses were investigated A) NLRP3 enhances ATM activation, B) NLRP3 inhibits ATM deactivation.

### Caspase-3/7 assay

Cells were cultured in 96-well plates and treated with 50 μM of Etoposide for 12 h or with 200 ng/mL Trail (Peprotech) and 1 mM MG132 (Sigma). Caspase 3/7 activity was assessed using the Caspase-Glo 3/7 assay reagent (Promega) following the manufacturer’s instructions. The luminescence was measured using a TECAN Infinite M200PRO luminometer microplate reader. To normalize the results, a second plate was stained with crystal violet and analyzed as described in the *crystal violet cytotoxicity assay* below.

### Crystal Violet cytotoxicity assay

Cells were stained with 0.5% crystal violet (Sigma-Aldrich Corp.) in 30% methanol for 20 minutes at room temperature. Cells were lysed in a 1% SDS (Sigma-Aldrich Corp.) solution. The absorbance of the solution was measured using a TECAN Infinite M200PRO microplate reader at a wavelength of 595 nm.

### Tissues from NSCLC patients

Frozen lung tumor tissues from non-treated patients were obtained from the Biological Resource Centre in Grenoble (Centre de Ressources biologiques de Grenoble) n°BB-0033-00069 and in accordance with the ethical laws. RNA was extracted from regions containing mainly malignant cells using Allprep RNA/DNA kit from Qiagen.

### TCGA data analysis

RNAseqV2 data of lung adenocarcinoma (LUAD) and corresponding clinical data were downloaded from The Cancer Genome Atlas TCGA data portal. Cox regression model was used to estimate hazard ratio (HR) and 95% confidence intervals (CIs) for overall survival (OS) and progression-free interval (PFI). Survival curve was estimated by the Kaplan–Meier method. Optimal cutoff for NLRP3 expression was chosen based on a maximally selected rank statistic^43^.

### Statistical analysis

Statistical analysis of the experimental data was performed using Graphpad. Unpaired group comparisons were done using two-tailed Student t-test for most figures, Mann-Whitney test for Figure 5A and B and multiple comparisons for two-way ANOVA for Figure 5C.

## RESULTS

### Loss of NLRP3 impairs γH2AX and P-ATM IRIF formation

To identify novel NLRP3 functions potentially linked to the regulation of the ATM pathway, we exposed HBEC3-KT cells to IR (2 Gy) and assessed the activation of the DDR pathway in the presence or absence of NLRP3. In control cells, the number of nuclear γH2AX (Ser139) IRIF peaked around 1 h after IR treatment and then decreased with time as the cells underwent DNA repair (Fig. 1A and Suppl. Fig. 1A). In contrast, in the absence of NLRP3 a significantly lower number of γH2AX IRIF were initially formed, and detected at all subsequent time points. This difference was greatest 1 h post IR and persisted over the time course of the study. ATM activation by DSBs relies on its monomerization which coincides with autophosphorylation on Ser1981 ^1^. Using the presence of P-ATM(Ser1981) IRIF as an endpoint for ATM activation we observed as early as 15 min post IR, fewer P-ATM foci in the absence of NLRP3, and the difference remained significant at 1 h and 5 h post-irradiation (Fig. 1B). Next, we tested whether the positive amplification loop required for optimal ATM activation was altered in the absence of NLRP3 by assessing the recruitment of MDC1 to DSBs. In siNLRP3-treated cells a decreased number of MDC1 foci were found compared with siCTL-treated cells, clearly illustrating a defect in the recruitment of MDC1 to DSBs, and, as a consequence, a defect in the ability to fully activate ATM (Fig. 1C and Suppl. Fig. 1B). 53BP1 is a DNA repair protein that also forms IRIF in response to DSBs but in an ATM-independent manner ^18^. The levels of 53BP1 foci were similar in the absence or presence of NLRP3 (Fig. 1D and Suppl. Fig. 1C), leading us to conclude that the decrease in the ability to form γH2AX and P-ATM foci supports a role for NLRP3 in the radiation-induced ATM DSB signaling pathway. This hypothesis was further investigated using mechanistic mathematical modeling (see Methods). Based on our experimental findings that NLRP3 KD reduced the observed number of P-ATM molecules recruited at DSBs, two hypotheses were investigated: A) NLRP3 enhances ATM activation, B) NLRP3 inhibits ATM deactivation (Suppl. Fig. 1D). Hypothesis A achieved a nearly perfect fit to data whereas hypothesis B was not able to reproduce the correct dynamics (SSR_A=0.052, SSR_B=0.29, Figure 1E). The model allowed to predict that the ATM activation rate in siNLRP3 conditions *k_IR_* was nearly half of that in siCTL cells as the ratio 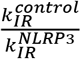 was equal to 1.66. Thus, the model A, which best recapitulated our data, supports the notion that NLRP3 enhances ATM activation.

### NLRP3 is required for optimal ATM activation

To determine whether NLRP3 played a broader role in the activation of the ATM pathway, we examined the phosphorylation of ATM downstream effectors that regulate cellular outcome in response to different DSB inducers. First, we showed that the absence of NLRP3 resulted in lower levels of KAP1 Ser824 phosphorylation in response to IR (Fig. 2A). Second, we tested the response of HBEC3-KT cells to two chemotherapeutic agents known to induce DSBs, etoposide (Eto), a topoisomerase II inhibitor, and hydroxyurea (HU), that depletes cellular dNTP pools, initially inducing stalled replication forks and, following longer exposure times, fork collapse and DSBs ^19^. Similarly, the absence of NLRP3 led to lower phosphorylation levels of KAP1 and p53 2 h to 6 h post-treatment with etoposide (Fig. 2B and C) or HU treatment (Suppl. Fig. 2A). NLRP3 down-regulation also resulted in decreased P-ATM irrespective of Eto concentrations (Suppl. Fig. 2B and C), and decreased γH2AX foci formation upon Eto treatment (Suppl. Fig. 2D). These results suggest that ATM is defective in the absence of NLRP3. Previous studies reported that ATM dysfunction induces ROS production in cells ^17,20^. Consistent with these studies NLRP3-depleted cells displayed enhanced ROS content compared to control cells, which did not increase further by the addition of the ATM inhibitor KU55933 (Suppl Fig. 2E). Collectively, these results strongly support the notion that NLRP3 is required for optimal ATM activation in response to DSBs.

**Figure 2.**
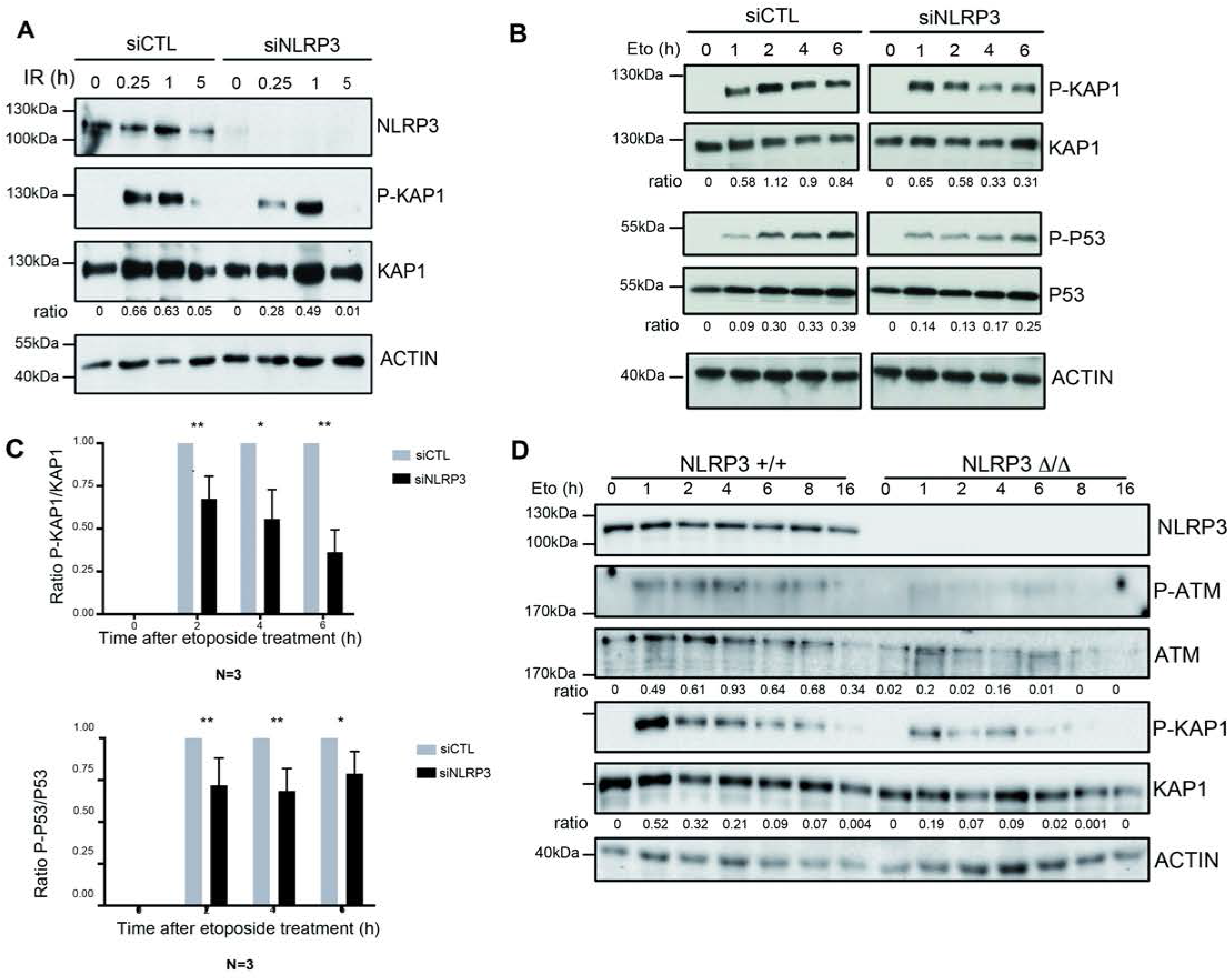
NLRP3 is instrumental for optimal ATM activation. (**A**) HBEC3-KT transfected with indicated siRNA were IR treated (10 Gy) and collected at different time points and P-KAP1 analyzed by immunoblotting. (**B**) HBEC3-KT control siRNA and NLRP3 siRNA were exposed to Eto (100 μM) for indicated time points and P-KAP1 and P-p53 were analyzed by immunoblot. (**C**) Relative quantification of immunoblot of 3 independent experiments as shown in (**D**). (**E**) Bone marrow-derived macrophages of wild type NLRP3 or NLRP3-depleted mice were treated with Eto 100 μM over time and P-ATM and P-KAP1 were analyzed by immunoblotting (representative of 2 experiments). Data represent mean ± SEM, *** P < 0.001, **** P < 0.0001, ns: not significant (unpaired t-test).

We then wondered whether NLRP3 was globally required for ATM activation using a heterologous model. Murine bone marrow-derived macrophages (BMDMs) either WT or NLRP3^Δ/Δ^ treated with Eto displayed deficient ATM activation, as evidenced by reduced ATM and KAP1 phosphorylation compared to controls (Fig. 2D). As we found A549 and H292 lung cancer cells did not express NLRP3, we re-expressed NLRP3 using a doxycycline-inducible system in these NSCLC tumor cell lines, and evaluated ATM activation after IR exposure by assessing the number of P-ATM and γH2AX IRIF. NLRP3 re-expression increased the levels of H2AX and ATM activating phosphorylations in both cell lines 1 h post-treatment (Fig. 3 A to D and Suppl. Fig. 3A to D). In addition, upon Eto treatment, the phosphorylation kinetics of the downstream effectors p53 and KAP1 in H292 re-expressing NLRP3 resembled those observed in non-tumoral control cells (Fig. 3E). Similar results were obtained in H520 IR-treated cells reconstituted with NLRP3 (Suppl. Fig. 3E). Thus, NLRP3 re-expression improved ATM activation.

**Figure 3.**
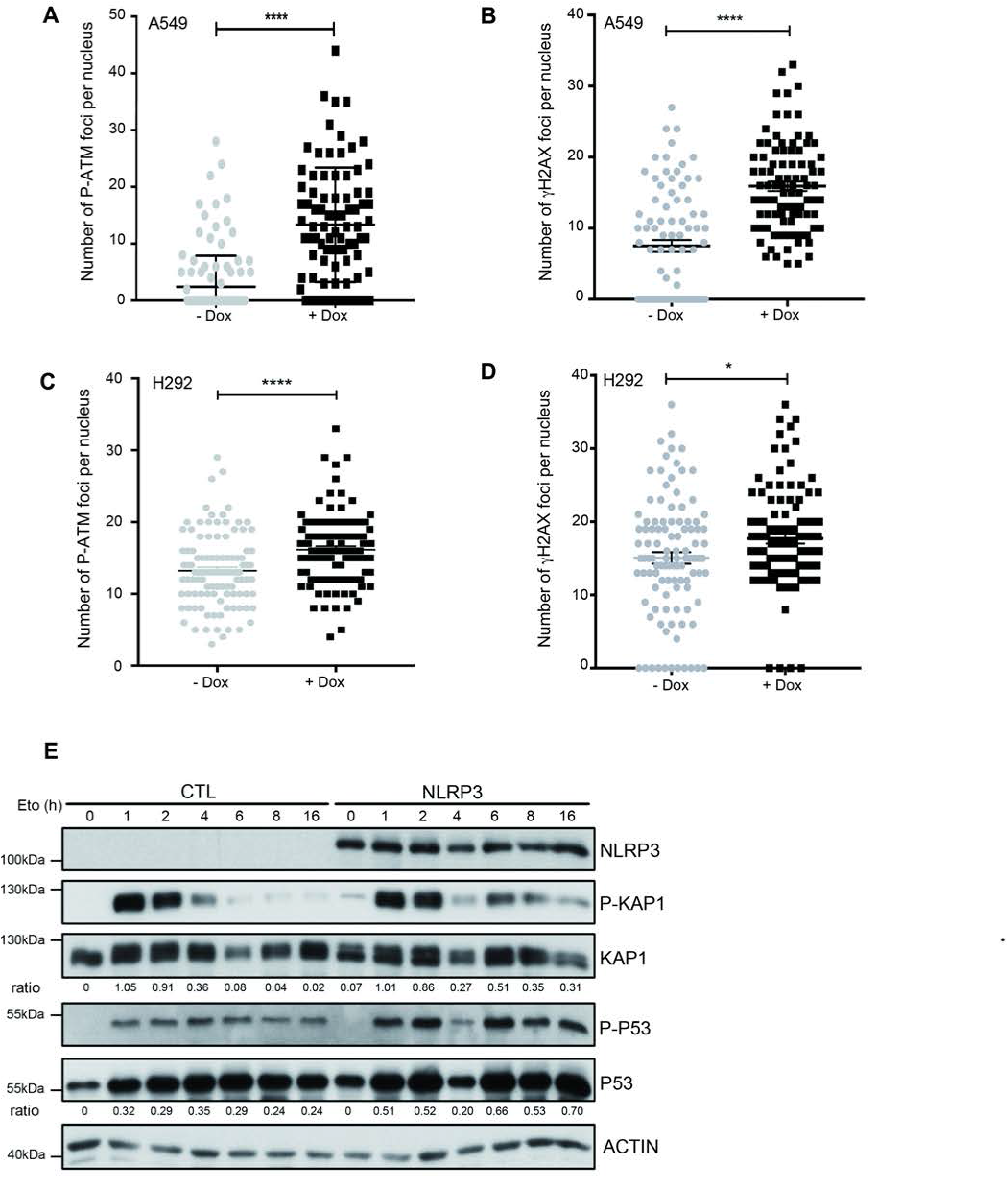
Expression of NLRP3 in NSCLC cell lines improves ATM activation after the induction of DNA DSBs. A549 (**A** and **B**) or H292 (**C** and **D**) cells stably expressing a doxycycline-inducible NLRP3 lentiviral vector (pSLIK NLRP3) induced or not with 0.5 μg/mL doxycycline were irradiated with 2 Gy and P-ATM (**A** and **C**) and γH2AX (**B** and **D**) IRIF assessed 1 h post-treatment. (**E**) H292 cells stably expressing the control or NLRP3 lentiviral vector were induced with doxycycline 24 h before being treated with etoposide over a time course of 16 h. KAP1 and p53 phosphorylation was analyzed by immunoblot at the indicated time points. One representative experiment out of 3. **** P < 0.0001, *** P < 0.001, * P < 0.05 (unpaired t-test).

### NLRP3 controls the ATM pathway independently of its inflammasome activity

To assess if this new role for NLRP3 in DNA damage signaling is dependent on its well-known inflammasome function, caspase-1 was knocked down using siRNA. The loss of caspase-1 did not alter the level of γH2AX after Eto treatment (Fig. 4A). In addition, no significant difference in the phosphorylation of H2AX was observed in HBEC3-KT cells treated with the pan-caspase inhibitor z-VAD-fmk or the caspase-1 inhibitor z-YVAD-fmk prior to HU exposure. These results would suggest that neither caspase-1 nor another caspase activity was involved in the activation of the ATM pathway, thus excluding apoptosis as a source of H2AX phosphorylation (Fig. 4B). In addition, DNA damage did not activate the inflammasome as IL-1β release was barely detectable in cell supernatants after HU or IR treatments, nor could we detect cleaved caspase-1 after Eto treatment (Fig. 4C, D, and E). Thus, DNA DSBs do not activate the catalytic activity of the inflammasome.

**Figure 4.**
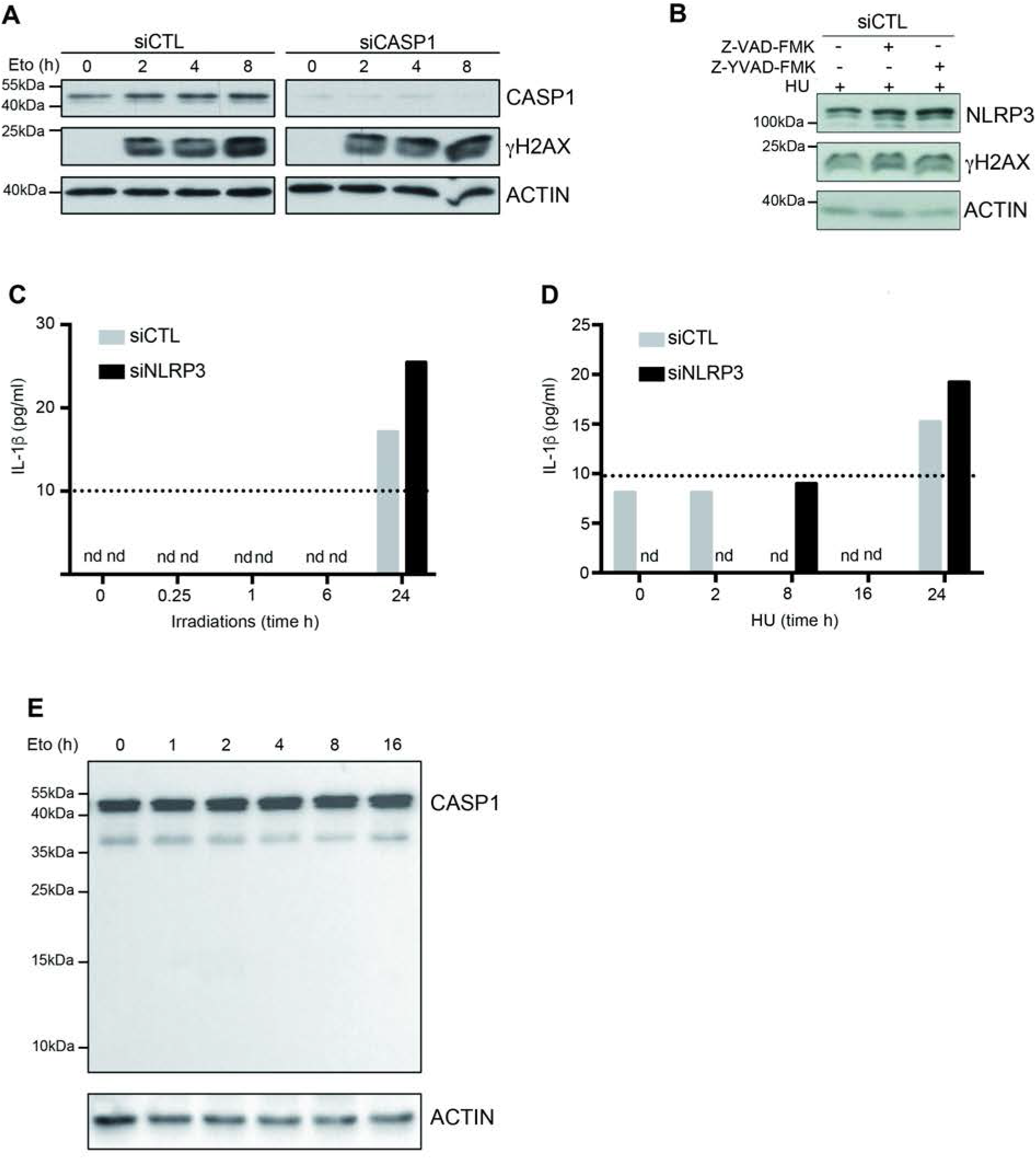
NLRP3 controls the DDR in an inflammasome-independent manner. (**A**) HBEC3-KT control siRNA and caspase-1 siRNA (siCASP1) were treated with Eto 100 μM and H2AX phosphorylation was monitored by immunoblot. (**B**) HBEC3-KT control siRNA were treated with the pan caspase inhibitor z-VAD-fmk (50 μM) or the caspase-1 inhibitor z-YVAD-fmk (50 μM) 30 min before HU treatment (2 mM, 16 h) and H2AX phosphorylation was analyzed by immunoblotting. (**C**) HBEC3-KT transfected with control or NLRP3 siRNA were treated with IR 2 Gy or (**D**) HU 2 mM for the indicated times and IL-1β was quantified in cell supernatants using a Luminex assay. The line indicates the detection limit. (**E**) HBEC3-KT cells were treated with Eto 100 μM over time and caspase-1 cleavage was analyzed by immunoblot. Actin was used as loading control. These data are from one representative experiment out of two independent experiments. n.d.: not detected

### NLRP3 forms a complex with ATM

To determine how NLRP3 regulates ATM activity, we tested whether these two proteins could interact with each other. In HeLa cells, that do not express endogenous NLRP3, co-immunoprecipitation of Flag-ATM pulled downed mCherry-NLRP3 (Fig. 5A). Moreover, Flag-NLRP3 co-immunoprecipitated endogenous ATM (Fig. 5B) ^21^. Eto treatment dissociated the interaction, suggesting that ATM and NLRP3 formed a complex under basal cell condition (Fig. 5B). We then mapped the NLRP3 domain involved in its binding to ATM. We showed that ATM interacted with the NACHT (domain present in neuronal apoptosis inhibitor protein, the major histocompatibility complex class II transactivator, HET-E and TPI), and the LRR (Leucin Rich Repeats) domains but not with the PYD (Pyrin) domain, which is known to mediate homotypic interactions involved in inflammasome formation (Fig. 5C) ^10^. NEK7 was used as a positive assay control, as it is a known partner of NLRP3 (Fig. 5C). We next investigated whether NLRP3 was able to translocate to the nucleus. Cell fractionation experiments revealed that endogenous NLRP3 was present in both the cytosolic and nuclear fractions (Suppl. Fig. 4A) supporting earlier findings ^22^. Interestingly, IF labeling of NLRP3 domains revealed nuclear localizations for the NACHT and the LRR domains, and a cytosolic localization for the PYD domain, which self-oligomerized as previously described for ASC PYD (Suppl. Fig. 4B) ^23^. Using this technique, short and full length (FL) NLRP3 were weakly detected in the nucleus (Suppl. Fig. 4B). However, live-imaging of mCherry-NLRP3 revealed its presence in the cell nucleus (Suppl. Fig. 4C). Consistently, co-immunoprecipitation experiments revealed that NLRP3 bound to IPO5 and XPO2 two proteins involved in nuclear import and export, respectively, which we identified by mass spectrometry (Fig. 5C) ^24,25^.

**Figure 5.**
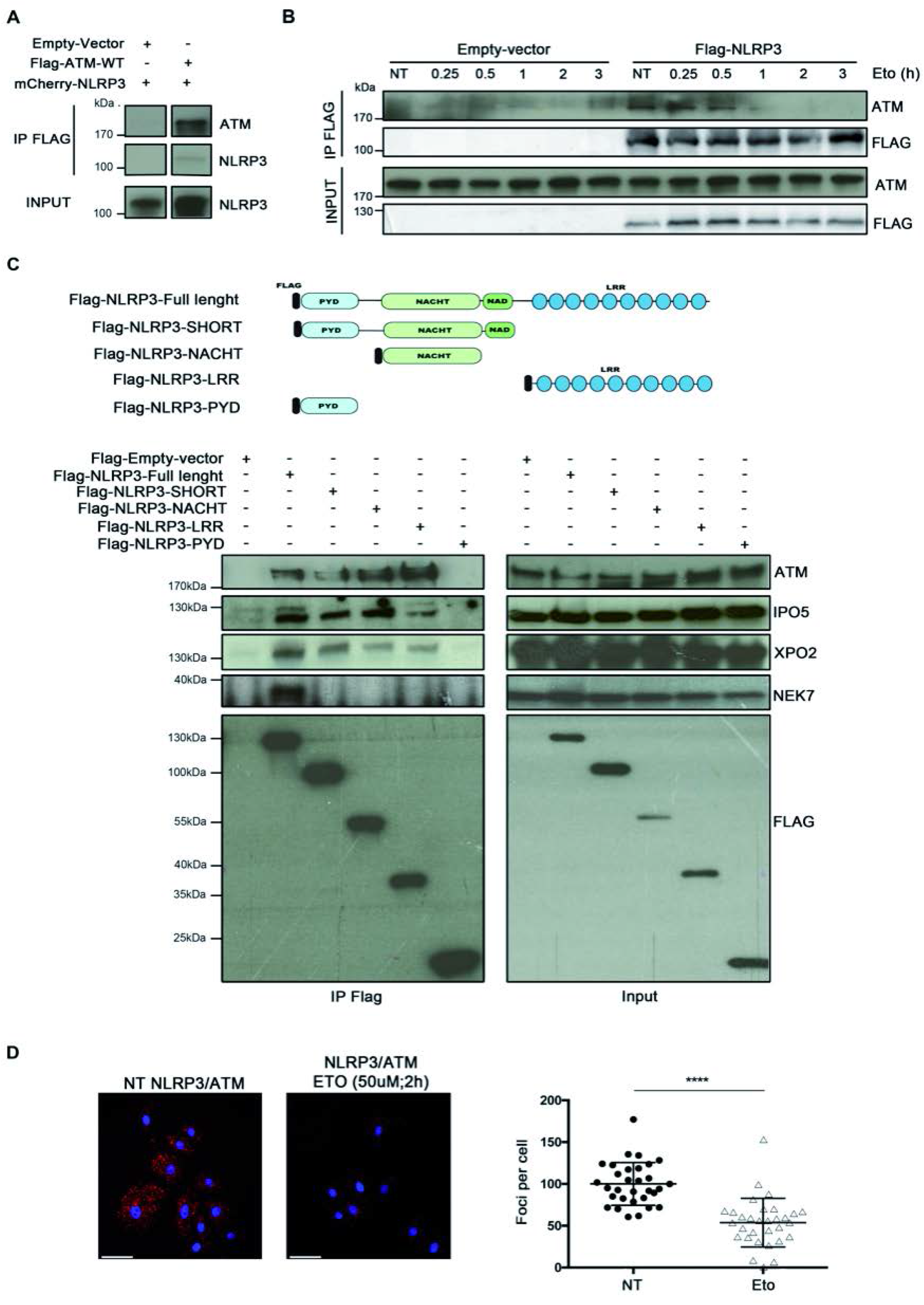
NLRP3 forms a complex with ATM. (**A**) mcherry-NLRP3 and Flag-ATM co-immunoprecipitate in HeLa cells. Immunoprecipitates (IP) and input were analyzed by immunoblotting. (**B**) HeLa cells expressing Flag-NLRP3 were treated or not with Eto for indicated time points. Co-immunoprecipitation of endogenous ATM was analyzed by immunoblotting. (**C**) Different flag-tag NLRP3 domain constructs were transfected in HeLa cells, and Flag-proteins were immunoprecipitated and pull-downed proteins were analyzed by immunoblotting. (**D**) Proximity Ligation Assay was performed in HBEC3-KT cells treated or not with Eto using anti-ATM and anti-NLRP3 (x40). Hoechst (blue) was used to stain nuclei. Scale bars 50 μm Signal quantification is shown on the graph on the right panel. NT: not treated.

Using Proximity Ligation Assay (PLA), we also showed in HeLa cells that Flag-NLRP3 and endogenous ATM formed a complex (Suppl. Fig. 4D). Importantly, we validated that endogenous ATM and NLRP3 interacted in HBEC3-KT cells, and that the complex was dissociated upon Eto treatment (Fig. 5D). We also observed after DNA damage that a smaller fraction of ATM was detected by IF in the nucleus of NLRP3-depleted cells compared with control cells (Suppl. Fig. 5A, B). Collectively, these results establish that under homeostatic conditions NLRP3 forms a complex with ATM, which dissociates upon DSB formation.

### NLRP3-depleted cells are resistant to genotoxic stress-induced cell death

Because ATM activity controls cell fate decisions in response to genotoxic stress, we next monitored cell death in response to Eto treatment. In NLRP3-depleted HBEC3-KT cells, less caspase-3/7 activity was detected compared with control conditions, and an increase in the number of viable cells was observed (Fig. 6A and B). These results suggest that decreased NLRP3 expression protects cells from etoposide-induced apoptosis. Indeed, the induction of *PUMA* and *NOXA/PMAIP1*, two p53 apoptosis effector genes, was significantly reduced in the absence of NLRP3 compared to control cells (Fig. 6C and D) ^26,27^. This response was specific to genotoxic stress as the induction of apoptosis via death receptor activation using a combination of TRAIL and MG132 did not result in impaired apoptosis in cells depleted for NLRP3 (Suppl. Fig. 6A and B).

**Figure 6.**
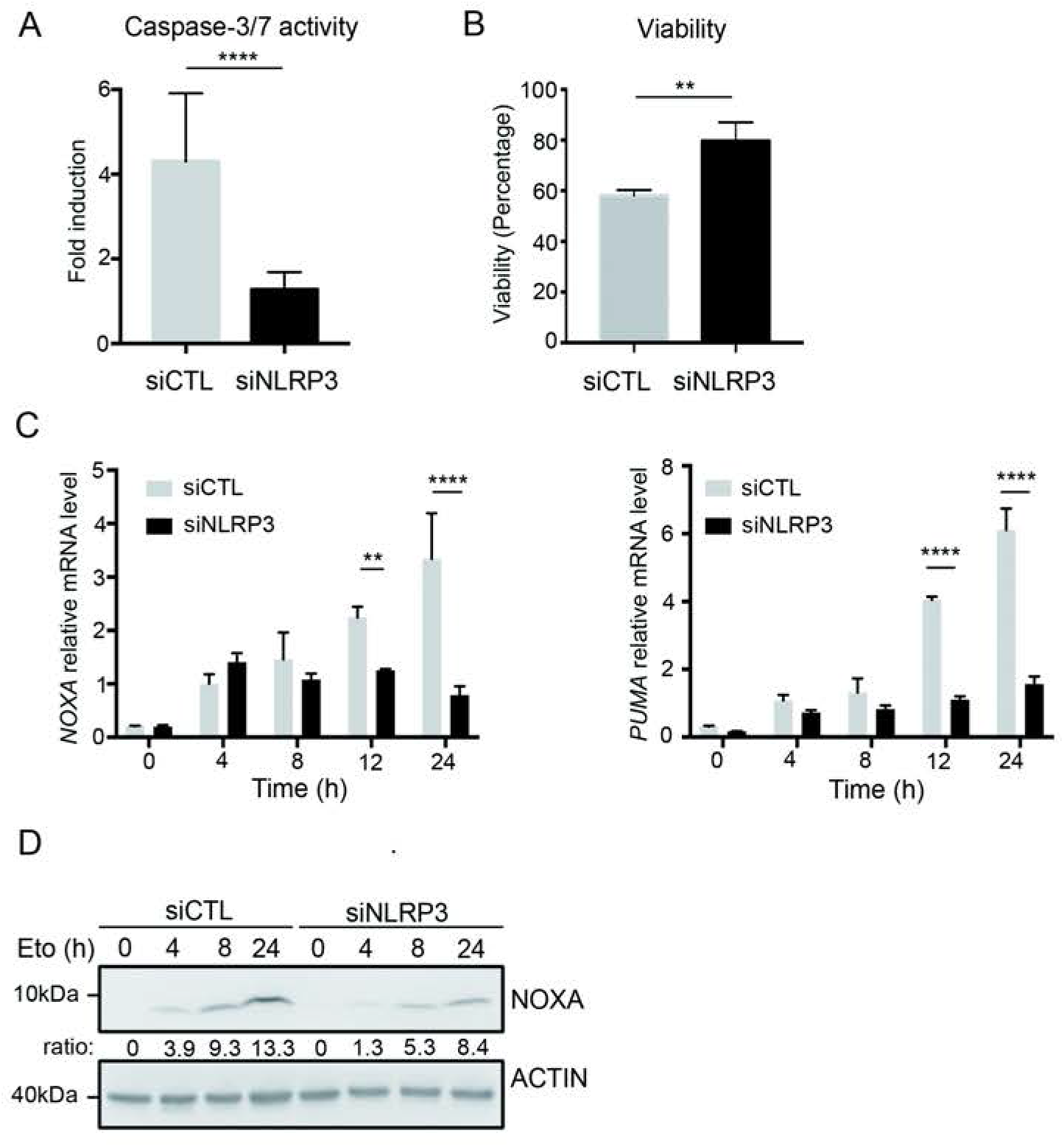
The absence of NLRP3 confers resistance to acute genotoxic stress. (**A** and **B**) HBEC3-KT transfected with the indicated siRNA were treated with Eto and (**A**) caspase-3/7 activity was then measured by luminometry and (**B**) cell survival by cristal violet cytotoxicty test. **** P < 0.0001, ** P = 0.0046 (unpaired t-test). Results are representative of three experiments. (**C**) *NOXA* and *PUMA* expression were assessed in HBEC3-KT treated with Eto at the indicated time points by Q-RT-PCR relative to *HPRT1* expression. **** P < 0.0001, ** P = 0.0035 (multiple comparisons for two-way ANOVA). (**D**) NOXA expression was assesed by immunoblotting. Actin was used as a loading control. Representative of two independent experiments.

### NLRP3 is down-regulated in NSCLC

GWAS studies reported that *NLRP3* is frequently mutated in NSCLC, but we found that A549 and H292 cells, isolated from NSCLC patients, do not express NLRP3. To extend this observation, we assembled a panel of NSCLC cell lines, and included 3 HBEC3-KT cell lines for comparison ^28^. Paradoxically, in most of the NSCLC cell lines, including cell lines reported to carry NLRP3 mutations, the NLRP3 protein was barely detectable (Fig. 7A and Suppl. Fig. 7A), and very low levels of *NLRP3* mRNA were observed by Q-RT-PCR in comparison with HBEC3-KT cells (Fig. 7B). Among the 3 HBEC3-KT lines, the HBEC3-ET cells did not express NLRP3, and those cells displayed properties of malignant transformation since they were able to grow in an anchorage-independent manner (Suppl Fig. 7B). These results suggest that NLRP3 expression is down-regulated in malignant cells. To validate these observations, we obtained a set of RNA samples from a cohort of patients with primary NSCLC and from adjacent normal lung tissues. As shown in Fig. 7C, *NLRP3* mRNA was detectable in normal lung tissues, while it was significantly down-regulated in NSCLC tissues. Finally, analysis of TCGA data showed that low NLRP3 expression in lung adenocarcinoma (LUAD) was associated with better overall survival and better progression-free interval (Fig. 7D and E). Altogether, these results suggest a down-regulation of NLRP3 in NSCLC, and a positive correlation in LUAD between NLRP3 levels and patient outcome.

**Figure 7.**
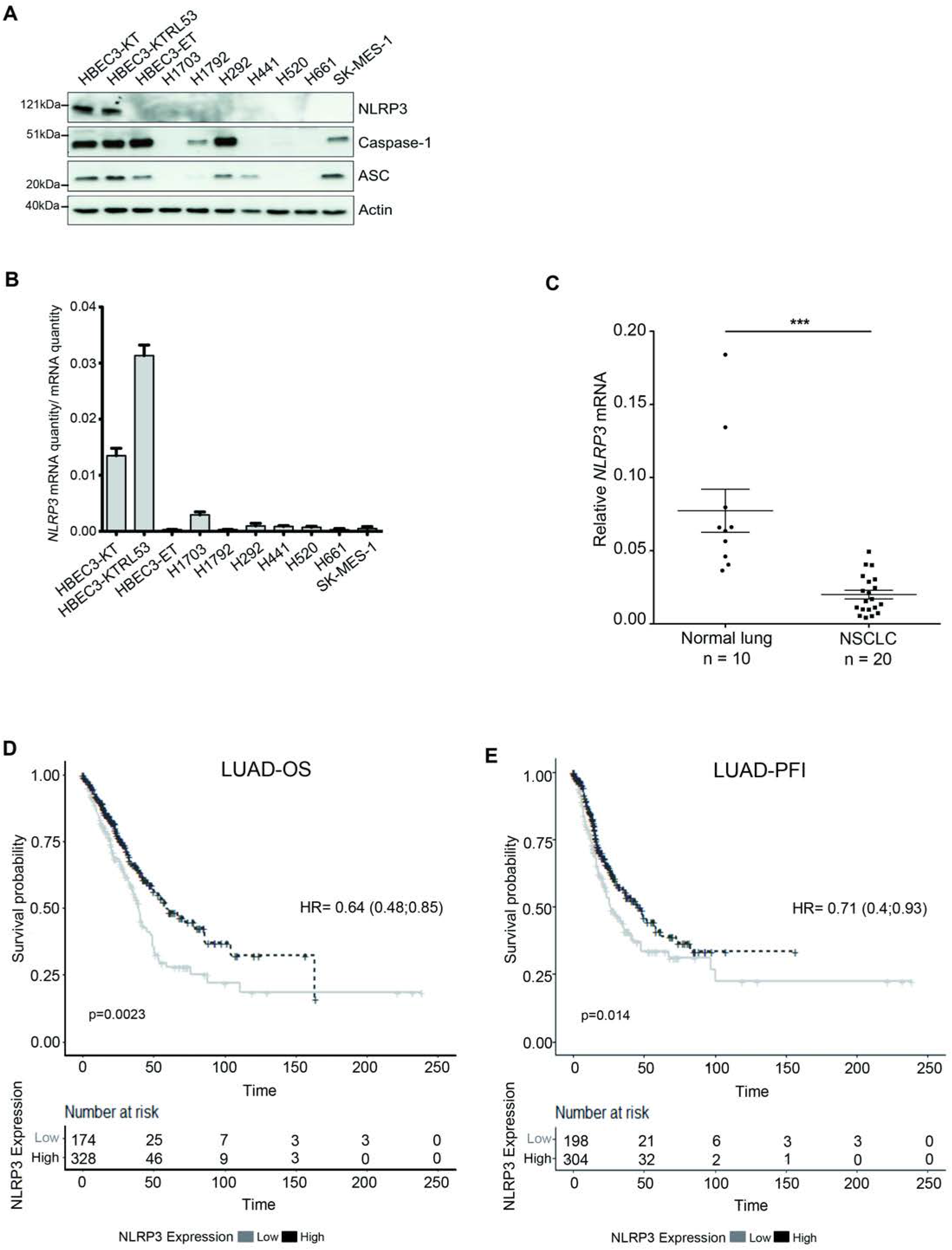
NLRP3 expression is reduced in human NSCLC compared to healthy tissue. (**A**) Protein levels of the NLRP3 inflammasome components NLRP3, caspase-1 and ASC were assessed by immunoblotting and (**B**) relative *NLRP3* mRNA by Q-RT-PCR in HBEC3-KT cells and in a panel of NSCLC cell lines. Results are representative of more than three experiments. (**C**) Relative *NLRP3* mRNA levels were determined by Q-RT-PCR in a cohort of non-treated primary tumors from NSCLC patients (n = 20) and the corresponding normal lung tissues (n = 10). Data represent mean ± SEM; *** P < 0.001 (t-test). Kaplan-Meier plots of patients overall survival (**D**) and progression free interval (**E**) in TCGA-LUAD dataset according to NLRP3 expression levels; patients were stratified according to the cutoff obtained from maximally selected rank statistic.

## Discussion

Functional links between innate immunity and DNA damage sensing pathways have been described. For instance, ATM was recently shown to be required in macrophages for optimal NLRP3 and NLRC4 inflammasome functions because its inactivation altered the ROS balance and therefore impaired inflammasome assembly ^17^. It was also suggested that DDR proteins such as RAD50 or BRCA1 are involved in the sensing of nucleic acids from viral pathogens in the cytosol ^29,30^. However, little is known about the contribution of PRRs to the sensing of stress like DNA damage. Here, we demonstrate that NLRP3 is crucial to reach optimal ATM activation. Under homeostatic conditions, our PLA data showed that NLRP3 forms a complex with ATM in the cytosol, suggesting that NLRP3 binds to the inactive ATM dimer. ATM has already been reported to be found in the cell cytosol ^20,31^. Upon DNA DSB formation, the complex dissociates to allow ATM relocalization and monomerization onto DNA breaks. In the absence of NLRP3, decreased levels of P-ATMSer1981 foci were observed herein, together with a decreased nuclear ATM pool. These observations suggest that NLRP3 may control either ATM translocation to the nucleus or ATM monomerization. Consequently, the formation of γH2AX and MDC1 IRIF, which are both essential for the positive ATM amplification loop signaling, were also impaired in NLRP3 KD cells ^6^. This led to a less active ATM as illustrated by the reduced phosphorylation of its substrates KAP-1 and p53, and, importantly, cells became more resistant to apoptosis.

No caspase-1 activation and no significant IL-1β production was detected upon DNA DSB induction. Our findings contrast with those of R. Flavell’s group, who demonstrated in mouse models that the severe damage caused to the gastrointestinal tract and the hematopoietic system in response to whole body γ-irradiation are due to the activation of the AIM2 inflammasome by DSBs, which cause massive cell death by caspase-1-dependent pyroptosis in these fast renewing tissues. These observations would suggest that tissue and species-specific differences may exist that clearly warrant further investigation ^32^.

Using different approaches which included restoring NLRP3 expression in NSCLC cell lines displaying low levels of inflammasome proteins, we identified a novel non-inflammasome function for NLRP3 in the DNA damage pathway. This previously unappreciated role for NLRP3 in the ATM pathway may be due to the fact that many common cellular models used in laboratories do no express NLRP3 (e.g. MEF, HeLa, 293, A549). Altogether, our results highlight that NLRP3 is not only a major player in innate immunity but is also a key factor involved in the preservation of genome integrity through the modulation of the ATM signaling pathway in response to DSBs.

The DDR is known to be a barrier to cancer in the early phases of tumorigenesis ^33,34^. *TP53* is frequently mutated in cancer, and the ATM pathway is down-regulated in many solid tumors: 11% of NSCLC carry somatic mutations in *ATM* and 41% of lung adenocarcinoma have reduced ATM protein expression ^35–38^. Although, several cancer genomic studies have reported that *NLRP3* is frequently mutated in NSCLC, our data actually suggest that in NSCLC primary human tissues and cell lines its expression is significantly lower compared to normal tissue. This down-regulation of *NLRP3* expression during malignant transformation may represent an additional mechanism to attenuate ATM and p53 signaling pathways, allowing cells to survive genotoxic stress, despite the presence of genome alterations. Thus, the loss of NLRP3, and the subsequent impairment of the ATM pathway could be an event allowing cells to progress towards malignancy.

## Acknowledgements

We thank John Minna for sharing the HBEC3-KT and NSCLC cells, Dr Foray’s team and Marine Malfroy for technical help and Agnès Tissier and Pascale Bertrand for helpful discussions. We thank Christophe Vanbelle and Christophe Chamot for their assistance on confocal microscope image acquisition. M.B. was supported by the ANRT, V.P. by the plan Cancer, Ligue Contre le Cancer Comité de l’Ain, the ARC foundation, and the Fondation pour la Recherche Médicale DEQ20170336744, A.L.H. by the ARC foundation and a Marie Skodolvska-Curie grant, N.G. was supported by the CLARA, and B.G. by the ARC foundation. B.P. was supported by the ERC.

## Author contributions

M.B.W., A.L.H., J.G., S.H., J.H. and V.P. designed and analyzed experiments. M.B.W., A.L.H., J.G., B.G., S.H., Y.C., L.G. and V.P. performed experiments. F.G., B.B., B.P., S.L., B.P. and N.G. provided reagents. M.B.W., A.L.H., J.G., S.H., J.H. and V.P. contributed to the manuscript writing and figure constructions.

---

**Supplementary Figure 1**. HBEC3-KT cells express a functional NLRP3 inflammasome.

(**A**) Immunoblot controlling the efficacy of the siRNA targeting NLRP3 (siNLRP3) against non-targeting siRNA (siCTL) of the HBEC3-KT irradiated cells. Actin was used as a loading control. (**B** and **C**) Representative pictures of HBEC3-KT cells transfected with control or NLRP3 siRNA and treated with 2 Gy for 1 h to assess MDC1 (**B**) or 53BP1 (**C**) foci formation was analyzed by IF. (x60), Hoechst (blue) was used to stain nuclei. Scale bars 10 μm. (**D**) Scheme displaying the hypothesis used for mathematical modeling of ATM and NLRP3 interactions showing the two hypotheses were investigated A) NLRP3 enhances ATM activation, B) NLRP3 inhibits ATM deactivation.

**Supplementary Figure 2**. ATM activity is impaired in the absence of NLRP3 in response to DNA damaging agents.

(**A**) HBEC3-KT siRNA-transfected cells were treated with 2 mM HU. At indicated time points, cells were lyzed and protein extracts analyzed by immunoblotting for NLRP3, γH2AX (S139), P-p53 (S15), P-KAP1 (S824). Actin was used as a loading control. (**B** to **D**) HBEC3-KT cells transfected with control or NLRP3 siRNA were treated with (**B**) 0.5 μM etoposide for 4 h and P-ATM foci were quantified, and (**C**) 100 μM Eto and mean fluorescence intensity was quantified in the nucleus, (**D**) 0.5 μM etoposide for 4 h and γH2AX foci quantified. (x60), Hoechst (blue) was used to stain nuclei. Data represent mean ± SEM; *** 0.001 < P (unpaired t-test). Scale bar 10 μm (**E**) ROS measurement was performed on HBEC3-KT cells transfected with control or NLRP3 siRNA using DCFDA probe in presence or in absence of 5 μM ATMi KU5593. One representative experiment out of 3.

**Supplementary Figure 3**. NLRP3 re-expression in tumoral cell lines facilitates ATM-dependent DNA damage signaling.

A549 (**A** and **B**) or H292 (**C** and **D**) stably expressing a doxycycline-inducible NLRP3 lentiviral vector treated or not with 0.5 μM of doxycycline were irradiated with 2 Gy for 1 h. Representative pictures of P-ATM (**A** and **C**) and γH2AX (**B** and **D**) IF staining that was quantified in Figure 3 A to D. (x60), Hoechst (blue) was used to stain nuclei. Scale bars 10 μm. (**E**) H520 cells stably expressing the NLRP3 or CTL vector were treated with 0.5 μg/mL of doxycycline 24 h before irradiation with 6 Gy. At indicated time points, cells were lyzed and protein extracts analyzed for NLRP3, γH2AX (S139), P-KAP1 and KAP1 by immunoblotting. Actin was used as a protein loading control.

**Supplementary Figure 4.** NLRP3 is localized in the cell cytosol and nucleus, but most NLRP3/ATM complexes are present in the cell cytosol.

(**A**) HBEC3-KT untreated (0) or irradiated (2 Gy) were separated and proteins from the cytosolic (C) and nuclear (N) fractions were analyzed by immunoblot. Tubulin was used as a marker of the cytosolic fraction and fibrillarin of the nuclear fraction. T is total lysate. (**B**) Confocal images illustrating the cellular localization of the different NLRP3 domains transfected in HeLa cells (x63). Hoechst (blue) was used to stain nuclei. (**C**) NLRP3 is detected in the cytosol and the nucleus compartment in live confocal image of mCherry-NLRP3 transfected in H292 cells. **(D)** Proximity Ligation Assay in HeLa cells transfected with an empty vector or a NLRP3-expressing vector using anti-ATM and anti-NLRP3 antibodies (x40). Hoechst (blue) was used to stain nuclei.

**Supplementary Figure 5**. NLRP3 silencing decreases nuclear ATM.

(**A** and **B**) HBEC3-KT sh control cells or knocked down for NLRP3 were left untreated (**A**) or irradiated at 2 Gy for 1 h (**B**). Total ATM was stained for immunofluorescence and the mean fluorescent intensity was quantified. Results are representative of two independent experiments (x60). Hoechst (blue) was used to stain nuclei. Scale bar 10 μm.

**Supplementary Figure 6**. NLRP3 does not control the extrinsic apoptosis pathway.

(**A**) HBEC3-KT cells transfected with control or NLRP3 siRNA were treated with Trail 200 ng/mL and MG132 1 mM for 12 h to induce death receptor-mediated apoptosis. Data represent mean ± SEM; ns: not significant (t-test). (**B**) Model of resistance to genotoxic stress caused by reduced NLRP3 expression. DNA DSBs activate the ATM kinase which phosphorylates many protein substrates involved in the control of the outcome to genotoxic stress. Down-regulation of NLRP3 impairs ATM activation resulting in decreased levels of phosphorylation of several ATM substrates, including p53, thus promoting cell survival.

**Supplementary Figure 7.** NLRP3 expression is reduced in human NSCLC compared to non-tumoral cells.

(**A**) NLRP3 inflammasome components, namely NLRP3, caspase-1 and ASC were assessed by immunoblotting in NSCLC cell lines. GADPH was used as a protein loading control. (**B**) Anchorage-independent growth ability was assessed in HBEC3-KT cell lines.

